# Facultative symbiont virulence determines horizontal transmission rate without host strain specificity

**DOI:** 10.1101/2023.02.16.528903

**Authors:** Suegene Noh, Emily R. Larson, Rachel M. Covitz, Anna Chen, Prachee R. Mazumder, Ron F. Peck, Marisa C. Hamilton, Robert A. Dettmann

## Abstract

In facultative symbioses, only a fraction of hosts are associated with a symbiont. Understanding why specific host and symbiont strains are associated can inform us of the evolutionary forces affecting facultative symbioses. Possibilities include ongoing host-symbiont coevolution driven by reciprocal selection, or priority effects that are neutral in respect to the host-symbiont interaction. We hypothesized that ongoing host-symbiont coevolution would lead to higher fitness estimates for naturally co-occurring (native) host and symbiont combinations compared to nonnative combinations. We used the *Dictyostelium discoideum* – *Paraburkholderia bonniea* system to test this hypothesis. *P. bonniea* features a reduced genome size relative to another *Paraburkholderia* symbiont of *D. discoideum*, indicating a significant history of coevolution with its host. Facultative symbionts may experience continued genome reduction if coevolution is ongoing, or their genome size may have reached a stable state if the symbiosis has also stabilized. Our work demonstrates that ongoing coevolution is unlikely for *D. discoideum* and *P. bonniea.* The system instead represents a stable facultative symbiosis. Specifically associated host and symbiont strains in this system are the result of priority effects, and presently unassociated hosts are simply uncolonized. We find evidence for a virulence-transmission trade-off without host strain specificity, and identify candidate virulence factors in the genomes of *P. bonniea* strains that may contribute to variation in benevolence.

**Lay summary:** Symbiotic relationships between hosts and their microbial partners are prolonged and intimate associations. Some of these relationships are obligatory for both a host and symbiont to survive, while others are facultative and each partner can survive without the other. In the latter case, some host individuals may be associated with a symbiont while others are not. Specific host and symbiont combinations can be the result of reciprocal adaptation between host and symbiont partners so that naturally co-occurring combinations are best suited for each other in terms of their biological fitness. On the other hand, the symbiont that a host is associated with may simply be the symbiont that arrived first, in what is called a priority effect. We sought to determine which possibility best explained naturally co-occurring combinations of host and symbiont strains of the social amoeba *Dictyostelium discoideum* and its symbiont *Paraburkholderia bonniea*. Our work demonstrates that *D. discoideum* and *P. bonniea* are in a stable facultative relationship. Specifically associated host and symbiont combinations are the result of priority effects, and *D. discoideum* hosts without symbionts are simply uncolonized. This work fills a gap in our understanding of the evolutionary forces affecting facultative symbiotic relationships. We also show for the first time that *P. bonniea* symbionts can spread among amoeba hosts when they aggregate together during the social stage of their life cycle.

## Introduction

Host-symbiont associations range from obligate or facultative depending on the degree of host-symbiont dependency (Sachs et al. 2011; Fisher et al. 2017). For obligate symbioses, when host and symbiont need each other to survive, we expect significant coadaptation between host and symbiont over a longer period of coevolution (Law and Dieckmann 1998). In contrast, for facultative symbioses in which host and symbiont can each survive in a free-living state, we expect coadaptation to a lesser degree or relationships to be more recent (Lo et al. 2016). Significant coadaptation in obligate symbioses, especially those that feature strict vertical transmission of symbionts, often result in symbionts with highly reduced genome sizes (Moran et al. 2008; McCutcheon and Moran 2012). In facultative symbioses, many of which include horizontal modes of symbiont transmission, symbiont genomes tend to be intermediate in size (Toft and Andersson 2010; Sachs et al. 2011; Lo et al. 2016; Fisher et al. 2017). While ongoing coevolution can lead to continued genome reduction for facultative symbionts, genome size may have reached a stable state if the facultative symbiosis has also stabilized. For example, *Glomeribacter* and Glomeromycota fungi are an example of an ancient facultative symbiosis supported by phylogenetic codivergence patterns (Mondo et al. 2012).

Many aspects of facultative symbioses are relatively poorly understood, including which factors determine distribution patterns of facultative symbionts among individuals of a host species (Niepoth et al. 2018). In many facultative symbioses, only a fraction of hosts are associated with a symbiont. Understanding why specific individual host and symbiont strains are associated with each other can inform us of the forces contributing to the evolution of facultative symbioses. Possibilities include ongoing host-symbiont coevolution driven by reciprocal selection, or priority effects that are neutral in respect to the host-symbiont interaction itself (Ganesan et al. 2022). Mutualistic coevolution could lead to specific combinations of host-symbiont strains that enhance each other’s fitness relative to other combinations (Rafaluk-Mohr et al. 2018). In contrast, priority effects in a symbiosis context have to do with the order of symbiont colonization and are typically considered when multiple symbiont species or strains coexist within the same environment as the host (Ganesan et al. 2022). If we extend priority effects to unassociated individual hosts in a facultative symbiosis, presently unassociated individuals may simply be uncolonized. In this sense, the question of what drives host-symbiont association patterns within a facultative symbiosis is related to questions of how host-microbe associations originate (Sieber et al. 2021) and how microbial communities are assembled (Tucker and Fukami 2014).

The potential for, and outcome of, long-term association between a host and any potential symbionts is determined by intrinsic factors such as host and symbiont genetics, as well as various extrinsic environmental factors (Schmid-Hempel 2011). For this work, we focused on the intrinsic factors of host defense and symbiont virulence and their potential contributions to fitness consequences of host-symbiont interactions. Host resistance (the ability to limit the extent of an infection) and tolerance (the ability to tolerate the physiological consequences of an infection) are the two large categories of host defense components that affect evolutionary outcomes for symbioses (Simms and Triplett 1994; Råberg et al. 2007; Ayres and Schneider 2012). The equivalent symbiont virulence components are proliferation (the ability to proliferate to a certain extent in an infection) and benevolence (the ability to cause beneficial or harmful physiological consequences of an infection) (Wollein Waldetoft et al. 2020). Given a specific combination of host and symbiont, variable host resistance and symbiont proliferation will lead to variation in symbiont density, while variable host tolerance and symbiont benevolence will lead to variation in host fitness consequences relative to symbiont density (Figure 1). Differentiating components of host defense or symbiont virulence is important because of their downstream evolutionary effects. For example, host resistance often limits the spread of infections while host tolerance can do the opposite and cause infections to increase in frequency in a population (Roy and Kirchner 2000; Råberg 2014). Targeting pathogen benevolence rather than proliferation may be more likely to prevent counter adaptation such as the evolution of antimicrobial resistance (Wollein Waldetoft et al. 2020).

**Figure 1.**
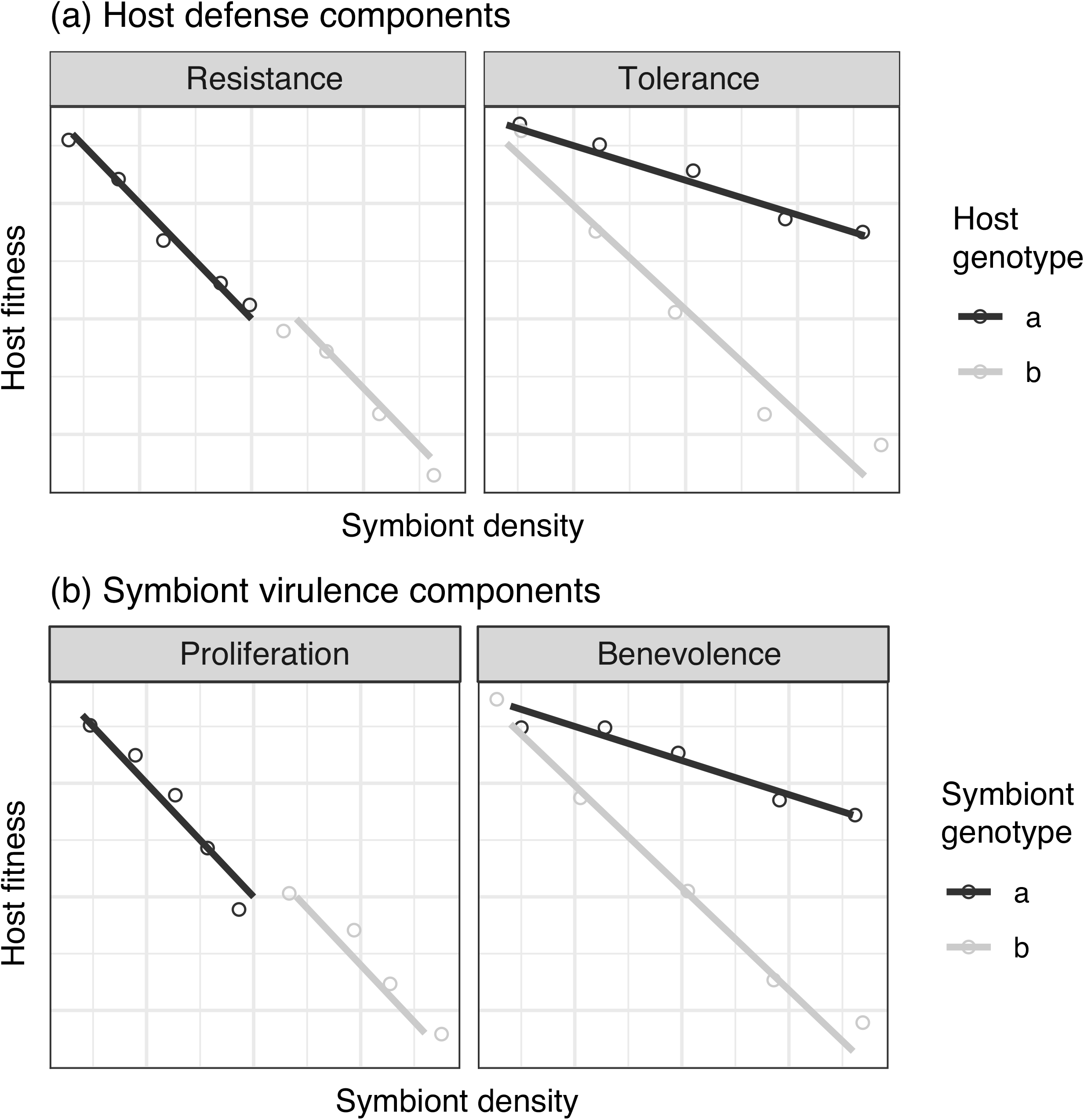
Theoretical differences between resistance-based and tolerance-based host defense, and proliferation-based and benevolence-based symbiont virulence. Both types of variation can be detected by measuring host fitness and detecting when (a) different host genotypes vary in responses to infection by the same symbiont, or (b) the same host genotype varies in responses to infection by different symbiont genotypes. (a) Hosts may vary in **resistance** to the same symbiont. More resistant hosts (a – black line; lower average symbiont density) will have overall higher fitness compared to less resistant hosts (b – gray line; higher average symbiont density). But the fitness cost per increase in symbiont density is similar for both hosts, as indicated by the similar slopes. On the other hand, hosts may vary in **tolerance** to the same symbiont. More tolerant hosts (a – black line; shallower slope) may have a similar average symbiont density to less tolerant hosts (b – gray line; steeper slope), but will suffer a lower fitness cost per increase in symbiont density. (b) The same host may suffer different fitness effects due to symbionts that vary in **proliferation**. The better proliferating symbiont (b – gray line; higher average symbiont density) will impart higher fitness costs to the host than the worse proliferating symbiont (a – black line; lower average symbiont density). But the fitness cost to the host per increase in symbiont density is similar for both symbionts. The same host may be infected with symbionts that vary in **benevolence**. More benevolent symbionts (a – black line; shallower slope) will impart a lower fitness cost per increase in symbiont density compared to more malevolent symbionts (b – gray line; steeper slope), though both symbionts may reach similar average densities. (Framework adapted from Raberg et al 2007 and Wollein Waldetoft *et al*., 2020.)

We used the *Dictyostelium discoideum* – *Paraburkholderia bonniea* system to determine whether ongoing host-symbiont coevolution affects which host-symbiont strain pairs naturally co-occur (are “native”) within a facultative symbiosis, and whether host defense and symbiont virulence contribute. The amoeba *D. discoideum* is an established model for understanding intracellular pathogen infections (Steinert and Heuner 2005). It forms a facultative symbiosis with three species of *Paraburkholderia*: *P. agricolaris*, *P. bonniea*, and *P. hayleyella* (DiSalvo et al. 2015; Brock et al. 2020). Roughly one quarter of soil-isolated *D. discoideum* strains carry one or occasionally multiple species of these symbiotic *Paraburkholderia* (Haselkorn et al. 2019). The three symbionts also provide an opportunity to contrast the effect of different evolutionary histories of association. Comparative genomics suggest that *P. bonniea* and *P. hayleyella* have evolved genomes that are half the size of other *Paraburkholderia* in the amoeba host environment (Noh et al. 2022). Patterns of host-symbiont coadaptation were previously tested for *P. agricolaris* and *P. hayleyella*, indicating that reciprocal adaptation is ongoing for native *P. hayleyella* hosts and symbionts. Native *P. hayleyella* hosts had a fitness advantage over nonnative hosts when cured and reinfected with *P. hayleyella*, while native *P. agricolaris* hosts did not have a similar advantage for *P. agricolaris* infections (Shu et al. 2018). Inside *D. discoideum* hosts, *P. hayleyella* grew faster than on its own while *P. agricolaris* did not (Garcia et al. 2019).

We hypothesized that ongoing host-symbiont coevolution would lead to higher fitness estimates for native host and symbiont combinations compared to nonnative combinations. We predicted coadapted intrinsic host and symbiont factors would contribute to fitness differences in host-symbiont associations. Using the fact that *D. discoideum* form social groups and symbiont horizontal transmission should be facilitated by these social groups, we examined host and symbiont fitness at the group level to better consider the evolutionary consequences of variation in fitness (Alizon and Michalakis 2015). We expected to find evidence of ongoing coevolution for *D. discoideum* – *P. bonniea* based on previous results from *P. hayleyella*, the sister species to *P. bonniea*, that contrasted from *P. agricolaris*.

## Methods

*D. discoideum* has single-cell and multi-cell stages in its life cycle. Vegetative single-cell amoebas feed on bacteria until food density becomes low and starvation begins. Next, in their social stage, starving amoebas aggregate into a multicellular form that ultimately forms fruiting bodies. The amoebas that survive the social stage become spores contained within the fruiting body sorus. We used the vegetative stage to expose *D. discoideum* amoebas to *P. bonniea* symbionts, and used spore counts after an unmanipulated social stage as a measure of host fitness. We then used a subsequent social stage to expose uninfected amoebas to pre-infected amoebas, and estimated symbiont horizontal transmission after this manipulated social stage as an important aspect of symbiont fitness.

### Focal host and symbiont strains

We used three native host strains of *D. discoideum* (QS395, QS433, QS859), three nonnative *D. discoideum* strains (QS4, QS17, QS18), and three *P. bonniea* symbiont strains (bb395, bb433, bb859). All *D. discoideum* strains were previously isolated from Mountain Lake Biological Station in Virginia, USA. The native host strains had been cured of their symbionts using tetracycline and verified as symbiont-free using PCR. For each replicate, host (with food bacteria *Klebsiella pneumoniae*) and symbiont strains were grown from freezer stock on SM/5 plates (2 g glucose, 2 g BactoPeptone (Oxoid), 2 g yeast extract (Oxoid), 0.2Lg MgCl_2_, 1.9Lg KH_2_PO_4_, 1Lg K_2_HPO_4_ and 15 g agar per liter). KK2 buffer (2.2 g KH2PO4 monobasic and 0.7 g K2HPO4 dibasic per liter) was used throughout for handling bacteria, *D. discoideum* spores and amoebas.

### Host fitness

We estimated host fitness at a range of infection prevalence for each host-symbiont combination (6 host strains x 3 symbiont strains). We estimate host fitness at the social group level rather than at the individual amoeba host level. Instead of symbiont density (e.g. Figure 1) we estimate symbiont infection prevalence within a group of amoebas, and instead of host fitness as a direct measure of individual host mortality we estimate group productivity given a degree of symbiont prevalence. *Experiment*: For each combination, *D. discoideum* spores from freshly grown fruiting bodies were collected and deposited on SM/5 plates at a density of 2 x 10^5^ spores per plate and RFP-labeled symbionts at multiplicities of infection (MOI) of 0 (control), 0.6, 3, and 15. We made triplicate sample plates per combination and left these plates to fruit at 21°C for 5-7 days. We then collected all fruiting bodies and estimated the number of spores per sample by counting 50x diluted spores on a hemocytometer and multiplying these counts by the total volume of spores. *Analysis*: Our models examine the relationship between host fitness and infection prevalence (not MOI). We estimated infection prevalence (percent of RPF+ infected spores in a sample of 100,000) by running a sample of spores from each plate on a BD FACSCalibur flow cytometer (Becton, Dickinson and Company, Franklin Lakes, NJ). FCS files were imported into FlowJo v10.8.1 for analysis. For host fitness, we took the percent of spores from each host-symbiont combination relative to the uninfected control that was prepared at the same time. For example, host strain QS4 was infected with bb395 at MOI of 0 (control), 0.6, 3, and 15. Spore counts of each individual sample plate were divided by the average spore count of the three uninfected control plates.

### Symbiont transmission

We estimated symbiont horizontal transmission for each host-symbiont combination (6 host strains x 3 symbiont strains). We estimate symbiont transmission at the social group level rather than at the individual amoeba host level using the social stage of the *D. discoideum* life cycle. For symbiont transmission we estimate the rate of transmission that occurs within a social group of interacting hosts given a previous degree of symbiont infection prevalence, rather than transmission from one host to another. *Pre-infection*: *D. discoideum* spores from freshly grown fruiting bodies were collected and deposited on SM/5 plates at a density of 2 x 10^5 spores per plate with RFP-labeled symbionts at MOI of 0 (control), plus an additional 3-5 MOI ranging from 0.3 to 30. Once these plates had fruited and amoebas were pre-infected, we collected fresh spores and deposited them on SM/5 plates at densities of 1 x 10^5 and 2 x 10^5 spores per plate with only food bacteria. *Experiment*: When amoebas were at log-phase growth roughly 36 hours later, uninfected amoebas were dyed with CellTracker Green CMFDA (Invitrogen) dissolved in DMSO. Pre-infected amoebas (RFP+) were carried along with the dyed amoebas, only exposed to DMSO, and treated in the same way through room temperature incubation and washes. Afterward, dyed amoebas were combined with infected amoebas at 1:0 (dyed only negative control), 0:1 (infected only), and 1:1 (mixed) ratios and placed on nitrocellulose filters to aggregate and develop into fruiting bodies in triplicate. 5-7 days later, we collected all fruiting bodies and their spores from these filters. *Analysis*: Our models examine the relationship between symbiont transmission and infection prevalence (not MOI). We estimated infection prevalence from the infected only sample and horizontal transmission from the mixed sample as indicated by the co-occurrence of dye and infection in a spore. As above, we ran spore samples on a BD FACSCalibur flow cytometer, and imported FCS files into FlowJo v10.8.1 for analysis. We applied gates for spores, then dyed cells from the negative control, and infected cells from one of the highest MOI infected only samples. A logical (AND) gate was applied to the mixed sample to find the percent of spores that were previously uninfected dyed amoebas that were now positive for symbiont infection (Figure 2).

**Figure 2.**
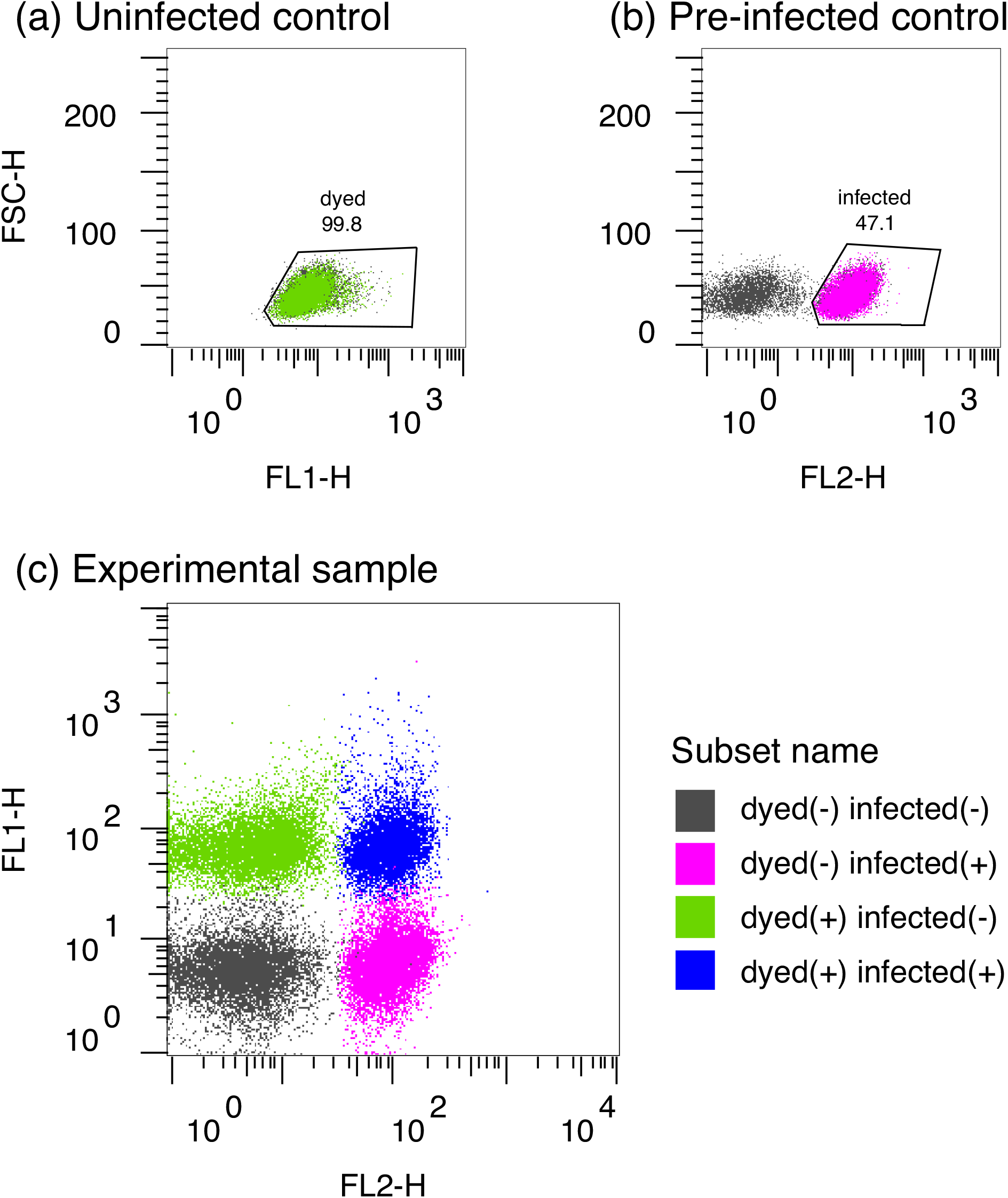
Representative samples from symbiont transmission experiment to demonstrate how symbiont transmission during a social stage is detected. (a) Dyed cells (green population) are gated on uninfected controls treated with Celltracker. (b) Infected cells with RFP-labeled symbionts (pink population) are gated on pre-infected controls. (c) Newly infected cells (blue population) are detected by applying logical gates to the experimental samples that mixed uninfected cells and pre-infected cells.

### Statistical analysis

Statistical analyses were performed in R v3.6.0 and with packages car v.3.0-12 (Fox and Weisberg 2019) and lme4 v.1.1-28 (Bates et al. 2015). For both host fitness and symbiont transmission, we fit linear models with mixed effects. For host fitness, we tested how percent spores (relative percent of spores produced) were affected by symbiont strain and host type (native or nonnative). We tested the effect of host type by coding the native combinations either strictly (e.g. only QS859 infected with bb859 would be considered native) or leniently (e.g. any native host of *P. bonniea* infected with any *P. bonniea* strain is considered native). We used infection prevalence as a continuous predictor and experiment date and host strain as random effects. For symbiont transmission, we fit a similar model but with percent transmission (percent newly infected spores that were previously uninfected amoebas) as the dependent variable. For all models we fit the most complex model first, then removed nonsignificant terms and compared the simpler model with the complex one using a chi-squared test. We examined effect sizes of factors in the final model using the package effectsize v.0.7.0 (Ben-Shachar et al. 2020) and its epsilon_squared() function. We also compared infection prevalence across the two experiments. We fit linear models with host strain as a random effect and the different number of social stages (nonnumeric factor; one for host fitness, two for symbiont transmission) and symbiont identity as fixed effects. We examined residuals of the final model for any indications that we had violated model assumptions. We used the package emmeans v.1.7.2 (Searle et al. 1980) and its emtrends() function for post-hoc tests of significant differences between pairwise slopes of infection prevalence by symbiont strain. We also used the emmeans() function for post-hoc tests of significant differences between means of infection prevalence by symbiont strain. *P*-values were adjusted using Tukey’s method.

### Genome analysis

We sequenced and assembled the genomes of bb395 and bb433 as follows. High molecular weight DNA was extracted using Lucigen MasterPure^TM^ Complete DNA and RNA purification kits (LGC, Teddington, UK). Extracted DNA was sent to University of Washington PacBio Sequencing Services for PacBio HiFi sequencing and MiGS for Oxford Nanopore (ONT) and illumina sequencing. Raw reads were cleaned using filtlong v0.2.1 (Wick 2023) and fastp v0.23.1 (Chen et al. 2018), then assembled and polished using Trycycler v0.5.3 (Wick et al. 2021) and Polypolish v0.5.0 (Wick and Holt 2022). Trycycler input files were created using Flye v2.9.1-b1780 (Kolmogorov et al. 2019) for both types of reads, Hifiasm v0.18.5-r499 (Cheng et al. 2022) for PacBio reads, and Raven v1.8.1 (Vaser and Šikić 2021) for ONT reads. Assembled contigs were re-oriented with Circulator v1.5.5 (Hunt et al. 2015). Genes were predicted using Prokka v1.14.6 (Seemann 2014). Pseudofinder v1.0 (Syberg-Olsen et al. 2022) was used to remove pseudogenes. Pan-genome analysis of the three *P. bonniea* genomes was performed using Roary v3.13.0 (Page et al. 2015). For examination of the differences between genomes, whole genomes were aligned using progressiveMauve (Darling et al. 2010) within Geneious Prime v2020.2.5 (https://www.geneious.com).

## Results

### Host fitness

We estimated the relationship between host fitness and symbiont infection prevalence by exposing native and nonnative *D. discoideum* hosts to three strains of *P. bonniea* at a range of multiplicities of infection. Native hosts were *D. discoideum* strains harboring their own *P. bonniea* symbiont strain when they were isolated from the wild. Host fitness decreased with increasing infection prevalence across all combinations of hosts and symbionts, but there was no difference between the native and nonnative host-symbiont combinations (Figure 3; Table 1). This was regardless of whether native status was defined using a strict or lenient definition (Supplementary material Table S1). Instead, symbiont strain identity significantly affected how host fitness responded to infection prevalence. For example, bb859 infections resulted in a significantly shallower slope in host fitness decline (β= -0.118, SE= 0.0172) compared to bb395 (β= -0.180, SE= 0.0158) and bb433 infections (β= -0.231, SE= 0.0157) (Figure 3). The effects of bb395 and bb433 infections on host fitness were not significantly different from each other. These results support a significant effect of symbiont benevolence variation on host fitness during initial infection. Based on our results, bb859 is more benevolent while bb395 and bb433 are more malevolent to all hosts.

**Figure 3.**
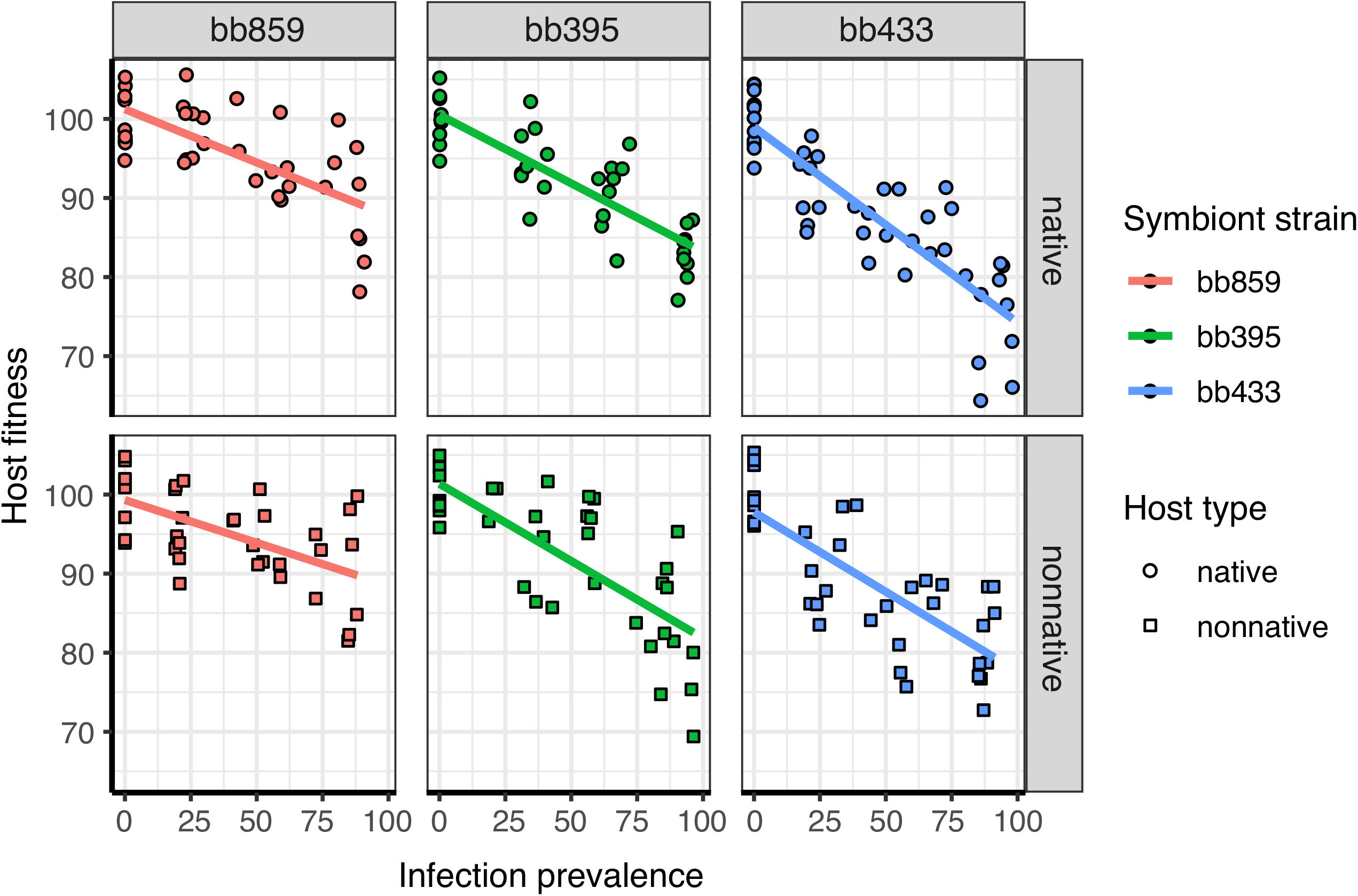
Host fitness was negatively correlated with infection prevalence and differed in slope but not average infection prevalence among symbiont strains after a single social stage. Infection prevalence was estimated per host-symbiont-MOI sample as the percent of infected spores that had RFP labeled symbionts in them. Host fitness was estimated by the percent of spores produced per host-symbiont-MOI sample, relative to uninfected hosts prepared at the same time as the infected host-symbiont combinations. There was no significant difference between native (top row; lenient definition) vs. nonnative (bottom row) host strains.

**Table 1.** Analysis of Deviance table for Host fitness with lenient native definition.

| **Variable** | **$\chi^2$** | ***df*** | ***P-value*** | **$\eta^2_p$** | **95% *CI*** |
| --- | --- | --- | --- | --- | --- |
| Infection prevalence | 376.002 | 1 | < 0.001 | 0.63 | (0.56, 1) |
| Symbiont strain | 17.998 | 2 | < 0.001 | - 0.06 | (0, 1) |
| Host type | < 0.001 | 1 | 0.981 | - 0.37 | (0, 1) |
| Infection prevalence :<br>Symbiont strain | 23.886 | 2 | < 0.001 | 0.09 | (0.04, 1) |

### Symbiont transmission

We estimated the relationship between symbiont transmission and symbiont infection prevalence by exposing uninfected amoebas to pre-infected amoebas during a manipulated social stage of the host life cycle. Horizontal transmission was detected when previously uninfected spores harbored *P. bonniea* at the end of this social stage. The degree of symbiont horizontal transmission increased with the infection prevalence of pre-infected amoebas that entered the social stage (Figure 4). This positive correlation confirms that the increase in infection is due to horizontal transmission rather than vertical transmission (Ebert 2013). This is consistent with previous reports that very few amoebas go through cell division during the social stage (Muramoto and Chubb 2008). As with host fitness above, native and nonnative host-symbiont combinations were no different in overall patterns of symbiont transmission (Table 2), regardless of how native status was defined (Supplementary material Table S2).

**Figure 4.**
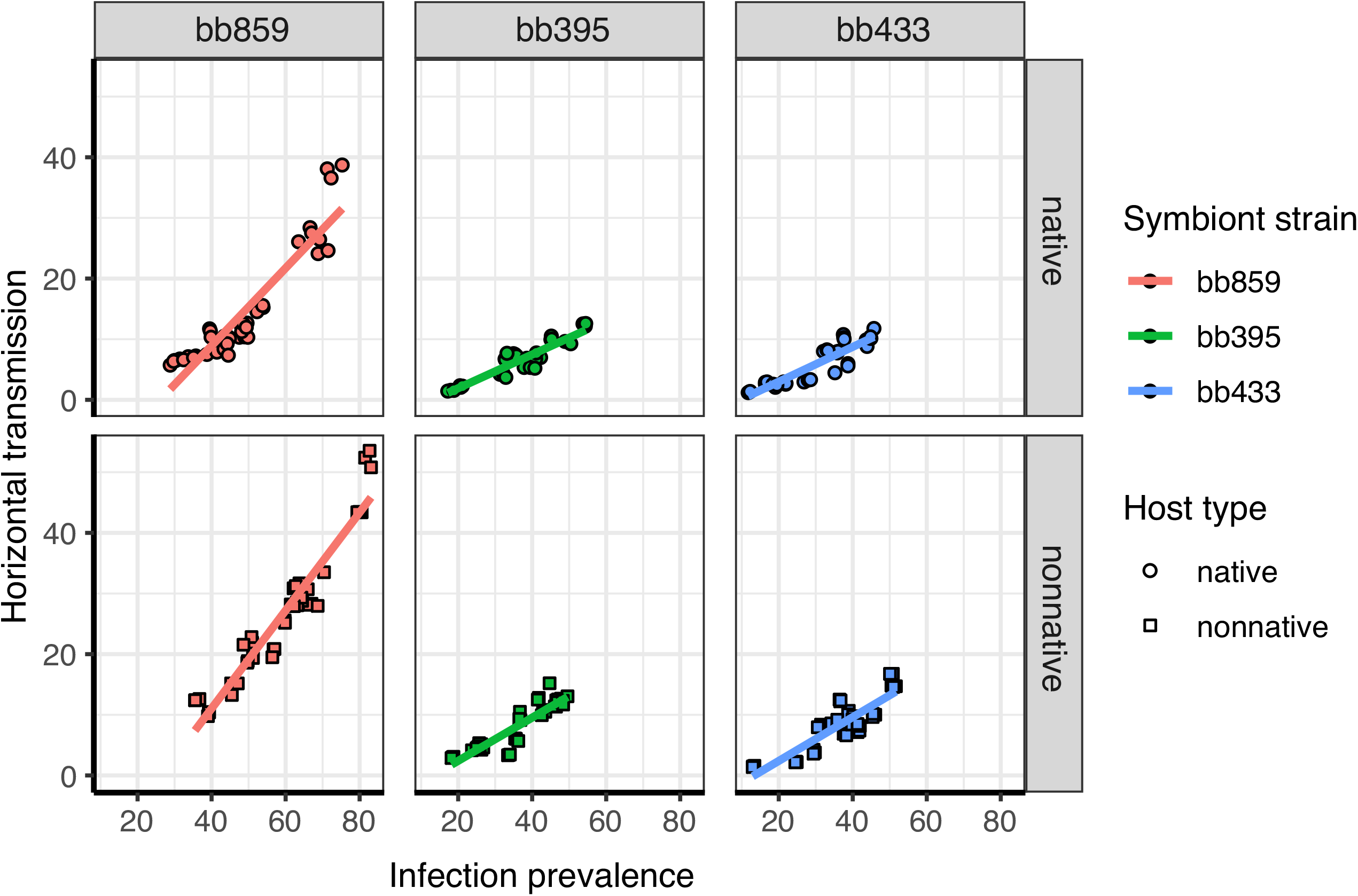
The rate of horizontal transmission was positively correlated with pre-infection prevalence and also differed in slope and average infection prevalence among symbiont strains after a second social stage. Infection prevalence was estimated per host-symbiont-MOI sample as the percent of RPF+ infected spores of the pre-infected control. Horizontal transmission was estimated by the percent of RFP+ infected spores in the test sample that were also positive for a green membrane dye. These dyed cells were uninfected prior to the experiment, during which they were combined with pre-infected (and undyed) cells. There was no significant difference in symbiont transmission among native (top row; lenient definition) and nonnative (bottom row) host strains.

**Table 2.** Analysis of Deviance table for Symbiont transmission with lenient native definition.

| **Variable** | **$\chi^2$** | ***df*** | ***P-value*** | **$\eta^2_p$** | **95% *CI*** |
| --- | --- | --- | --- | --- | --- |
| Infection prevalence | 484.965 | 1 | < 0.001 | 0.77 | (0.71, 1) |
| Symbiont strain | 149.430 | 2 | < 0.001 | 0.43 | (0.20, 1) |
| Host type | 2.092 | 1 | 0.148 | 0.53 | (0.25, 1) |
| Infection prevalence :<br>Symbiont strain | 109.289 | 2 | < 0.001 | 0.65 | (0.52, 1) |
| Infection prevalence :<br>Host type | 35.991 | 1 | < 0.001 | 0.30 | (0.17, 1) |
| Symbiont strain : Host<br>type | 9.509 | 2 | 0.009 | 0.14 | (0.01, 1) |

In this experiment, infected bb859 reached higher infection prevalence and showed a higher rate of horizontal transmission compared to bb395 and bb433 (Figure 4). Pre-infection was established in one social stage, and the experiment was performed in a subsequent social stage. The three *P. bonniea* strains used in this experiment do not significantly differ in their per amoeba-spore bacterial density (Miller et al. 2020). Therefore, higher infection prevalence in bb859 compared to the other strains does not necessarily indicate that bb859 is better at proliferation. However, bb859 symbionts transmitted at a significantly higher rate (β= 0.525, SE= 0.0244) beyond that simply due to bb859 infections occurring at higher infection prevalence compared to the other two strains bb395 (β= 0.361, SE= 0.0343) and bb433 (β= 0.249, SE= 0.0289) (Figure 4). When native hosts were leniently defined but not when strictly defined, infection prevalence and host type interacted to reveal that nonnative hosts became infected by bb859 at a slightly higher density compared to native hosts and therefore also transmitted symbionts at a higher level.

### Synthesis of host and symbiont fitness

We observed a significant difference in infection prevalence between the two experiments that is likely due to differences in symbiont strain-specific virulence (Table 3; Figure 5). The main difference between the design of the two experiments is that for *Host fitness* we used newly-infected amoebas, but for *Symbiont transmission* we used amoebas that had been infected in a previous vegetative stage, to minimize the potential for symbionts to be newly acquired from the environment and to maximize our ability to detect horizontal transmission among amoebas. In addition, we infected amoebas for the symbiont transmission experiment at a wider range of MOI (0.3-30) compared to the host fitness experiment (0.6-15) in order to achieve as wide a range of infection prevalence as possible. We found that we were unable to push the maximum of these ranges higher by using a higher MOI for the initial infection.

**Figure 5.**
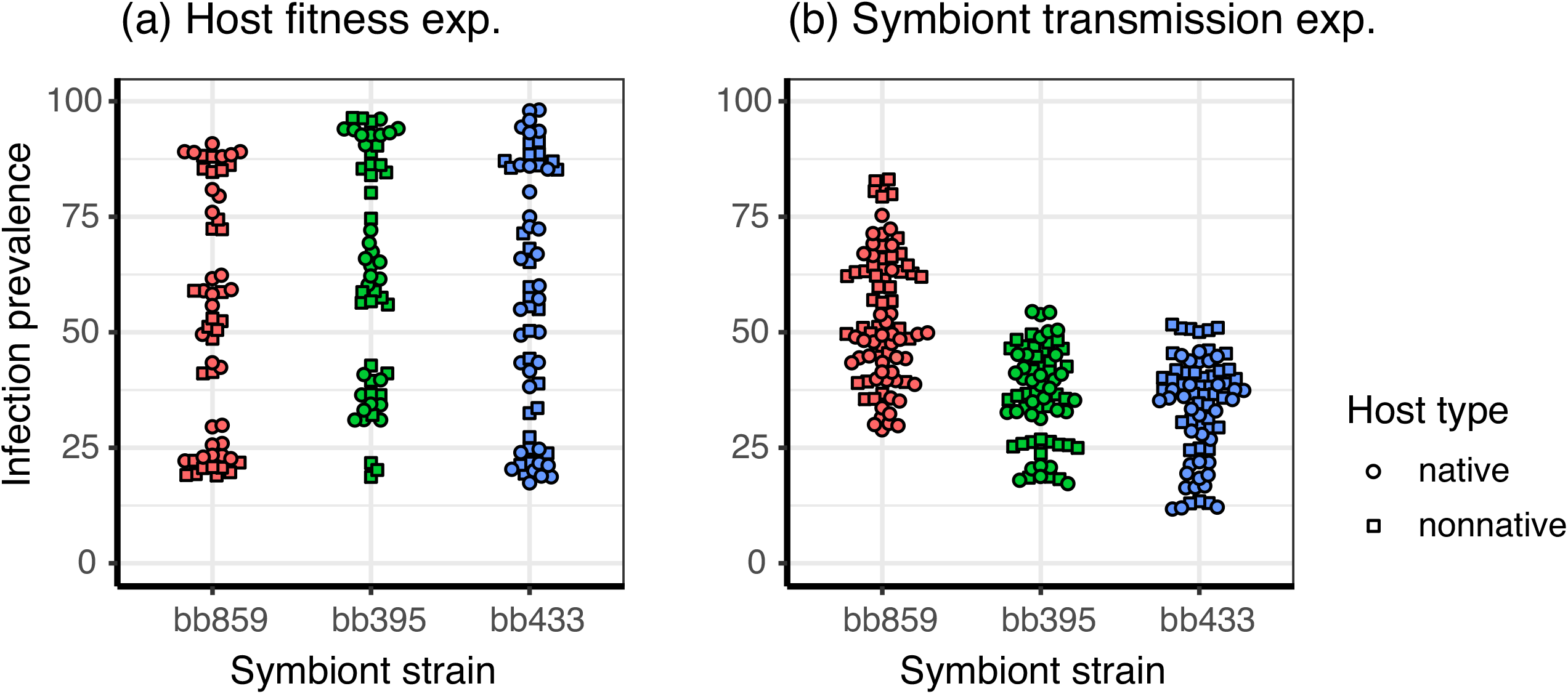
Infection prevalence was significantly different when amoebas were observed after one (a) vs. two social stages (b) after infection. More malevolent strains (bb395 and bb433) reach lower infection prevalence among amoeba hosts compared to more benevolent strains (bb859) over time.

**Table 3.** Analysis of Deviance table for infection prevalence across experiments.

| **Variable** | **$\chi^2$** | ***df*** | ***P-value*** | **$\eta^2_p$** | ***95% CI*** |
| --- | --- | --- | --- | --- | --- |
| Number of social stages | 71.196 | 1 | < 0.001 | 0.16 | (0.11, 1) |
| Symbiont strain | 17.789 | 2 | < 0.001 | 0.02 | (0.00, 1) |
| Number of social stages : Symbiont strain | 39.197 | 2 | < 0.001 | 0.09 | (0.05, 1) |

Combined observations from both experiments suggest that variation in symbiont benevolence leads to variation in symbiont prevalence among *P. bonniea* strains infecting amoeba hosts in the longer term. We observed no differences in infection prevalence after the first vegetative stage – social stage cycle (Figure 5(a)). But more malevolent strains bb395 (μ= 36.5, SE= 2.38) and bb433 (μ = 33.8, SE= 2.29) become significantly less prevalent among amoeba hosts compared to more benevolent strain bb859 (μ= 53.5, SE= 2.17) over the course of two vegetative stage - social stage cycles (Figure 5(b)). A higher rate of horizontal transmission in benevolent strains compared to malevolent strains during social stages contributes to this result (Figure 4).

### Candidate virulence factors

The three *P. bonniea* strains used in this experiment do not significantly differ in their per spore bacterial density (Miller et al. 2020). Yet the fact that we observe virulence variation in benevolence suggests that bb395 and bb433 possess malevolent virulence factors that bb859 lacks, or bb859 possesses one or more benevolent virulence factors that the other strains do not. Whole genome alignment revealed two structural variants each approximately 13 kilobases in size present in strains bb395 (1:108428 – 122213 and 1:2654181 – 2667432) and bb433 (1:108428 – 122213 and 1:2658228 – 2671479) and absent in bb859. These variants likely resulted from integration of plasmids as both regions have genes encoding putative plasmid recombinases near the 3’ end. We also identified an approximately 6 kilobase structural variant in strain bb859 (2:708878 – 715517) that lacks closely-related homologs in the other sequenced genomes of *Paraburkholderia* symbionts of *D. discoideum* (Brock et al. 2020).

## Discussion

The reduced-genome symbiont *P. bonniea* is able to persistently infect *D. discoideum* and form a facultative symbiosis in which only a fraction of hosts are associated with this symbiont (DiSalvo et al. 2015; Haselkorn et al. 2019; Miller et al. 2020). Based on our results, *D. discoideum* and *P. bonniea* are in a stable facultative symbiosis, where host-symbiont coevolution is primarily historical rather than ongoing. Host and symbiont strains responded similarly to association in native and nonnative combinations. We observed variation in virulence among *P. bonniea* strains but its effect did not vary for native and nonnative *D. discoideum* hosts.

### Strain level variation in symbiont virulence – virulence-transmission trade-off

Horizontal transmission of *Paraburkholderia* symbionts was previously assumed, but we demonstrate it for the first time in this study. In addition, we found evidence that strain level variation in symbiont virulence affects transmission by enabling higher rates of horizontal transmission in benevolent strains compared to malevolent strains. The virulence-transmission trade-off hypothesis posits that if there is a biological link between symbiont virulence and transmission (e.g. because symbionts use host resources to replicate), symbionts should evolve toward a level of virulence where transmission is greatest (Anderson and May 1982). This hypothesized trade-off has proven difficult to observe for several reasons, including the influences of host range, multiple symbiont infections, and host population structure that can affect host-symbiont interactions in complex ways (Alizon et al. 2009; Leggett et al. 2013). However, recent meta-analyses indicate that there is partial support for the trade-off hypothesis in several biological systems (Acevedo et al. 2019). The three core predictions of the trade-off hypothesis are as follows: (i) within-host symbiont replication rates have a positive relationship with symbiont virulence; (ii) within-host symbiont replication rates have a positive relationship with symbiont transmission rates; and (iii) symbiont virulence and transmission have a trade-off through the negative relationship between symbiont virulence and host recovery rates.

Because we estimated host fitness and symbiont transmission at the social group level rather than at the individual amoeba host level, our evidence that supports each of these points may be an indirect test of the virulence-transmission trade-off, as the original theory is formulated based on investigating symbiont density in individual hosts. However, considering fitness at the population level can facilitate our understanding of how host defense and symbiont virulence evolve (Alizon and Michalakis 2015). We find support for each of the core predictions of the virulence-transmission trade-off hypothesis. Symbiont infection prevalence had a negative relationship with host fitness (i; Figure 3), and symbiont infection prevalence had a positive relationship with symbiont horizontal transmission (ii; Figure 4). Lastly, variation in symbiont benevolence led to increased horizontal transmission of benevolent strains compared to malevolent strains (iii; Figure 4).

Transmission dynamics are an important missing piece of information in further understanding the evolution of the facultative symbiosis between *D. discoideum* and *Paraburkholderia*. Given the persistence of *Paraburkholderia* symbiont infections in the lab and the facultative aspect of the symbiosis itself, we expect these symbionts to transmit among hosts using both vertical and horizontal routes. We did not test for vertical transmission and specifically designed our experiment to use the social stage so that only horizontal transmission was possible. Our results clearly show the presence of significant horizontal symbiont transmission among *D. discoideum* amoebas in the social stage of their life cycle. The coexistence and relative influences of both types of transmission routes are better understood in other amoeba-bacteria symbioses (Herrera et al. 2020).

### Lack of variation in host defense – ecology and geography

Our results indicate that native hosts of *P. bonniea* do not possess enhanced host defenses to *P. bonniea* compared to nonnative hosts. These results contrast with evidence of host-symbiont coadaptation in *P. hayleyella* and its native hosts (Shu et al. 2018; Garcia et al. 2019). Both *P. hayleyella* and *P. bonniea* have reduced genomes and are sister species to each other (Brock et al. 2020). The lack of host strain-specific counteradaptation to *P. bonniea* virulence variation may be related to the ecology of host and symbiont. *P. bonniea* is the rarest of *Paraburkholderia* symbionts of *D. discoideum* (Haselkorn et al. 2019; DuBose et al. 2022). Rare encounters between new hosts and *P. bonniea* may not have a significant impact on *D. discoideum* populations, particularly if *D. discoideum* itself is sparsely distributed across soil landscapes. The fitness cost of infection for rare *P. bonniea* may be insufficient for hosts to evolve defenses against it (Anderson and May 1982).

Another potential reason why we did not find evidence of host defense variation may be because our native and nonnative hosts were collected from the same geographical locality. In other words, coevolution in this system may occur at the population level rather than at the strain level. Examples of strain-level coadaptation in facultative symbionts are limited. In pea aphids, coadaptation with their symbionts goes in both directions. Pea aphids from a population (*Lotus pedunculatus* biotype) that typically carry *Hamiltonella defensa* were more likely to establish symbiosis when newly infected with *Hamiltonella* compared to hosts from another population (*Lotus corniculatus* biotype) (Niepoth et al. 2018). However, in a tripartite host-symbiont-pathogen system, host-by-symbiont and symbiont-by-pathogen genotype by genotype interactions affected how protective *Regiella* symbionts were to pea aphids against the fungal pathogen *Pandora*, but not in an way that supported coadaptation between native host and symbiont genotype combinations (Parker et al. 2017). *Xenorhabdus* spp. are obligate symbionts to various *Steinernema* spp. nematodes. Evidence of host-symbiont coadaptation has been found in a couple different pairings, including between strains of *Xenorhabdus nematophila* and *Steinernema carpocapsae* (Chapuis et al. 2009), and between *Xenorhabdus bovienii* strains and their various *Steinernema* species hosts (Murfin et al. 2015).

Alternatively, the difference in results between *P. bonniea* and *P. hayleyella* may be due to previous experimental approaches that do not account for host or symbiont fitness as a function of infection prevalence. We plan to apply the experimental approach of the current study to the relationship between *D. discoideum* and *P. hayleyella*, which is known to cause more detrimental fitness consequences to nonnative hosts (Shu et al. 2018; Miller et al. 2020). If previous results hold for *P. hayleyella,* it would provide support for the evolution of reduced antagonism in *P. bonniea*. Amoeba hosts experience only a mild reduction in fitness to *P. bonniea* infection compared to *P. agricolaris* or *P. hayleyella* (Miller et al. 2020). Reduced antagonism is a potential outcome of symbiosis that is favored when virulence-transmission tradeoffs are present and new hosts are rare (Yamamura 1993; Johnson et al. 2021). Both of these conditions appear to hold for *P. bonniea* and support our interpretation that the *D. discoideum* – *P. bonniea* relationship is a stable facultative symbiosis.

### Candidate virulence factors for benevolence variation

We identified two structural variants that are present in the genomes of bb395 and bb433 but absent in bb859, as well as an additional structural variant that is present in the genome of bb859 but absent in the other two strains. We examined these regions for potential virulence factors that might contribute to the variation in benevolence we observed among *P. bonniea* strains.

Several candidate genes may confer increased malevolence to bb395 and bb433. Among these, PB395_00119/ PB433_00119 encode a putative member of the peptidase S8 family (also called subtilisin-related peptidases) that are known contributors to the pathogenesis of *Streptococcus pneumoniae* (Ali et al. 2021). Interestingly, there are no closely-related homologs in other sequenced *Paraburkholderia* genomes, but proteins with the highest similarity in Genbank are found in *Burkholderia pseudomallei* (MBF3536330.1, 93% identity over the length of the protein) and *Ralstonia solanacearum* (NKA33280.1, 93% identity over the length of the protein), which are respectively pathogens of mammals and plants. Potentially, this peptidase may be introduced into *D. discoideum* cells and modify host responses to infection.

In the other putative plasmid integration site, PB395_02359/ PB433_02360 is a member of the Xenobiotic Response Element (XRE) family of transcriptional regulators. It is most similar to a homolog in the opportunistic human pathogen *Burkholderia cepacia* complex (WP_060080935.1, 80% identity over the length of the protein). In these proteins, the putative DNA-binding domain is at the N terminus of the protein and the C-terminal portion of the protein contains a predicted peptidase domain. This architecture is similar to the AlpR gene in *Pseudomonas aeruginosa* which is involved in the pathway of programmed cell death and has been demonstrated as a virulence factor (McFarland et al. 2015).

One gene in particular appears to be an interesting candidate for a benevolence factor for strain bb859. PBONN_03430 encodes a putative member of a Type I secretion system known to secrete a wide variety of proteins from Gram negative bacteria (Spitz et al. 2019). An interesting possibility is that the PBONN_03403 protein may be part of a complex that secretes proteins that modulate activities in *D. discoideum* cells to prevent host cell death and allow bb859 to transmit to other host cells. We are actively working to assess the role of these genes in *P. bonniea* virulence.

### Conclusions

Our work demonstrates that ongoing coevolution is unlikely for *D. discoideum* and *P. bonniea.* The system instead represents a stable facultative symbiosis. Specifically associated host and symbiont strains in this system are the result of priority effects rather than ongoing host-symbiont coevolution, and presently unassociated hosts are simply uncolonized. In addition, we found evidence for a virulence-transmission trade-off without host strain specificity. A benevolent strain had a higher horizontal transmission rate compared to malevolent strains during the social stage of the host life cycle, regardless of host strain. Lastly, we identified candidate virulence factors in *P. bonniea* genomes, paving the way for future research in determinants of strain level variation in benevolence.

## Data availability

Full genome sequences for *P. bonniea* strains are available through NCBI Genomes (txid2152891). Strain bb859 (SAMN09651436) was submitted previously and bb395 and bb433 were newly submitted (SAMN35686367, SAMN35686368). Additional data and code supporting this work can be accessed at: https://github.com/noh-lab/social-infection

## Acknowledgements

We thank Susanne DiSalvo, who generously shared the RFP-labeled *P. bonniea* strains, and the Strassmann-Queller lab who generously shared the *D. discoideum* strains used in this work. We thank past and current members of the Noh lab (Caroline Lunt, Gus Shuster) who contributed moral support for this project during its development. Lily Khadempour, Laura Runyen-Janecky, Chris Moore, and Judy Stone provided helpful comments that improved previous drafts.

This work was supported by the National Institutes of Health and its National Institute of General Medical Sciences by an Institutional Development Award (IDeA) under Grant Number P20GM103423 (subaward to SN) and Colby College startup funds.

