## Supplementary material for "Facultative symbiont virulence determines horizontal transmission rate without host strain specificity"

Table S1. Analysis of Deviance table for Host fitness with strict native definition

| **Variable** | $\chi^2$ | ***df*** | ***P-value*** | $\eta^2_p$ | ***95% CI*** |
|---|---|---|---|---|---|
| Infection prevalence | 377.339 | 1 | < 0.001 | 0.63 | (0.57, 1) |
| Symbiont strain | 12.888 | 2 | 0.002 | - 0.08 | (0, 1) |
| Host type | 1.944 | 1 | 0.163 | < 0.001 | (0, 1) |
| Infection prevalence : Symbiont strain | 23.849 | 2 | < 0.001 | 0.09 | (0.04, 1) |

Table S2. Analysis of Deviance table for Symbiont transmission with strict native definition

| **Variable** | $\chi^2$ | ***df*** | ***P-value*** | $\eta^2_p$ | ***95% CI*** |
|---|---|---|---|---|---|
| Infection prevalence | 418.058 | 1 | < 0.001 | 0.60 | (0.53, 1) |
| Symbiont strain | 67.724 | 2 | < 0.001 | 0.65 | (0.00, 1) |
| Host type | 1.371 | 1 | 0.242 | 0.12 | (0.00, 1) |
| Infection prevalence : Symbiont strain | 85.187 | 2 | < 0.001 | 0.68 | (0.53, 1) |
